# Genetic and epigenetic influences modulating the tissue specific regulation of the cannabinoid receptor -1 gene (CB_1_); implications for cannabinoid pharmacogenetics

**DOI:** 10.1101/544585

**Authors:** Elizabeth A. Hay, Philip Cowie, Andrew McEwan, Dana Wilson, Ruth Ross, Perry Barrett, Roger G. Pertwee, Alasdair MacKenzie

## Abstract

Cannabinoid receptor-1 (CB_1_) represents a potential drug target against conditions that include obesity and substance abuse. However, drug trials targeting CB_1_ (encoded by the CNR1 gene) have been compromised by differences in patient response. Towards addressing the hypothesis that genetic and epigenetic changes within the regulatory regions controlling CNR1 expression contribute to these differences, we isolated the human CNR1 promotor (CNR1prom) and demonstrate its activity in primary cells and transgenic mice. We also provide evidence of CNR1prom in CB_1_ autoregulation and its repression by DNA-methylation. We further characterised a conserved regulatory sequence (ECR1) in CNR1 intron 2 that contained a polymorphism in linkage disequilibrium with disease associated SNPs. Deletion of ECR1 from mice using CRISPR genome editing significantly reduced CNR1 expression in the hippocampus. These mice also displayed reduced ethanol intake and hypothermia response to CB_1_ agonism. Moreover, human specific C-allele variants of ECR1 (ECR1(C)) drove higher levels of CNR1prom activity in hippocampal cells than did the ancestral T-allele. We further demonstrate a role for the AP-1 transcription factor in driving higher ECR1(C) activity. In the context of the known roles of CB_1_ the current study suggests a mechanism through which ECR1(C) may be neuroprotective in the hippocampus against stress. The cell-specific approaches used in our study to determine the functional effects of genetic and epigenetic changes on the activity of tissue-specific regulatory elements at the CNR1 locus represent an important step in gaining a mechanistic understanding of cannabinoid pharmacogenetics.

## Introduction

The cannabinoid-1 receptor (CB_1_) is expressed in areas of the nervous system that include the hypothalamus and the hippocampus where CB_1_ plays a critical role in appetite regulation (1) and neuroprotection against stress (2). For this reason, CB_1_ has been explored as a target for drugs to treat diseases including obesity and depression (1, 3). There are numerous examples of the successful use of cannabinoid drugs in the treatment of disease. For example, Sativex (a combination drug comprising of plant cannabinoids, delta-9-tetrahydrocannabidiol (THC) and cannabidiol (CBD)) supresses the chronic pain and spasticity associated with multiple sclerosis (4, 5). However, the synthetic CB_1_ antagonist rimonabant, marketed as an appetite suppressor, was withdrawn because 26% of patients reported depression, anxiety and feeling of suicidality (6). Moreover, there is strong evidence of a genetic component to the psychotic, cognitive and addictive side effects of CB_1_ agonists (7, 8). Given the potential benefits of the pharmacological manipulation of CB1 it is essential to gain a better understanding of cannabinoid pharmacogenetics to facilitate the development and application of safe and effective cannabinoid-based therapeutics.

The current study examines the hypothesis that polymorphic and epigenetic changes in the regulatory regions that ensure the correct tissue specific expression of the CNR1 gene (encodes the CB_1_ receptor), alter the regulation of CNR1 which may contribute to differences in response to drugs that target the CB_1_ receptor. Using a unique combination of comparative genomics, reporter assays in primary and transformed cell cultures, chromatin immunoprecipitation, in-vivo CRISPR/cas9 gene editing in mice, in-vivo gene expression and phenotype analysis we have dissected and characterised some of the components of the regulatory mechanisms that modulate the tissue specific expression of the human CNR1 gene. The current study also highlights the role of regulatory polymorphisms in the differential activation of these regulatory components by CB1 agonists and identifies a role for DNA methylation in controlling CNR1 gene expression. We will discuss these novel observations within the context of cannabinoid pharmacogenomics.

## Materials and Methods

### Plasmid constructs

#### pCNR1prom-Luc

The human genomic fragment containing the *CNR1* promoter (CNR1 prom) was amplified from human placental DNA using the following primers (9):

> CNR1prom for; 5’-GATAACCTTTTCTAACCACCCACCTAG-3’,
>
> CNR1prom rev. 5’-GCGGAAAAGAAGTGGAGAAG-3’

and cloned into the *EcoRI* and *SacI* restriction sites to replace the generic TATA box promoter of the pGL4.23 luciferase reporter construct to create the pCNR1prom-Luc (Fig 2A).

#### pECR1C/TLuc

The ECR1 enhancer sequence was also recovered from pGEM-T easy (Nicoll et al 2012) and the minor T allele of ECR1 was re-created using site-directed mutagenesis where the template DNA was a previously established pECR1(C)Gem-Teasy construct, primers for site-directed mutagenesis were:

> ECR1(C296T)For5’-ACTTCCTTTGAGAGTTCATTACTAATATGGCTTAGGCTTTGG-3’
>
> ECR1(C296T)Rev5’-CCAAAGCCTAAGCCATATTAGTAATGAACTCTCAAAGGAAGT-3’

Production of pECR1(C)CNR1promLuc and pECR1(T)CNR1promLuc firefly luciferase reporter constructs was achieved by cloning the ECR1(C) or ECR1(T) regions from the pGEM-Teasy parent constructs using *AatII* and *SalI* sites and ligating into *AatII* and *XhoI* sites of the pCNR1prom luciferase construct, placing ECR1(C) or ECR1(T) upstream of the *CNR1prom* fragment to form pECR1C/TLuc (Fig 2A)

*pCNR1prom-LacZ*. pCNR1prom was recovered from pCNR1promLuc and cloned into the NotI and SpeI sites of the LacZ reporter plasmid p1229 (10).

*pCNR1 promLucCpG free*. A CpG dinucleotide-free version of the CNR1prom-Luc reporter plasmid was created using the pCpGL-basic reporter plasmid (Klug and Rehli, 2006) and cloning the *CNR1prom* fragment from pCNR1promLuc reporter using *BglII* and *NcoI* sites.

### Plasmid methylation

Subsequently, 5μg of pCNR1promLucCpG free plasmid was methylated in a reaction containing 1x NEB buffer 2, 640μM SAM and 20U *M.Sss1* enzyme (New England Biolabs) at 37°C followed by inactivation (65°C, 20mins) at various time points up to 1 hour (Full methylation). Levels of methylation were monitored using digestion by the *HpaII* enzyme that is sensitive to methylated DNA (supplementary data) and methylated plasmids were transfected as described above.

### Primary Cell culture

1-day old male and female Sprague-Dawley neonate rats were euthanized in accordance with current UK Home Office schedule 1 guidelines. Hippocampal or hypothalamic tissues were dissected into ice cold Neurobasal-A medium (Life Technologies). Cells were dissociated using a combination of trypsin and papain solution followed by gentle agitation from pooled tissues to reduce variability and cultured in poly-d-lysine coated 24-well plates as previously described (11) at a density of 180,000 cells per well as assessed using a Biorad TC10 cell viability counter. Cells were maintained at 37°C in 5% CO_2_ for up to 7 days *in vitro* in Neurobasal-A medium supplemented with 2% B27, 2mM L-glutamine, 50 μg/ml Streptomycin and 50U Penicillin (Life Technologies) with the medium changed on the first day after plating and then every 3 days. Cultures were transfected 4 days after plating with luciferase plasmids (Figure 2A) plus renilla luciferase plasmid (pGL4.74) as a transfection normalisation control. Neuromag transfection reagent was used as per manufacturer’s instructions (OZ Biosciences). 24 hours after transfection cultures were treated with Win55,212-2 for 16 hours.

### Transformed cell culture and transfection

SH-SY5Y cells (94030304, ECAC, UK) were cultured in Dulbecco’s Modified Eagle Medium (DMEM; Gibco, UK) containing low Glucose (5.5mM), L-Glutamine (4mM), and Sodium Pyruvate (1mM). Medium was supplemented with 10% (v/v) heat-inactivated Foetal Bovine Serum (FBS; Gibco) and 1% (v/v) Penicillin-Streptomycin (Pen-Strep; Gibco, UK). Cells were co-transfected with luciferase reporter plasmids (Figure 2A) or a positive control (pAP1-3, Addgene 71258) and a renilla normalisation control (pGL4.74; Promega) together with pcDNA-FLAG-FosWT (Addgene 8966) or empty expression vector (pcDNA3.1, Thermofisher; V79020) plasmid using jetPRIME as per manufacturer’s instruction (Polyplus Illkirch-France).

### Luciferase reporter assays

All luciferase reporter assays were performed either 24 (SH-SY5Y cells) or 48 hours (Primary cells) after transfection as per manufacturer’s instructions (Promega, UK). Luciferase expression was quantified using Dual Luciferase Reporter assay system and a GloMax 96 microplate luminometer.

### Chromatin immunoprecipitation (ChIP)

was performed from primary hippocampal neurons transfected with either the ECR1(C) or ECR1(T) reporter constructs (1.2μg/5×10^6^ cells in 10cm plates) as described above and incubated for 48 hours. Following cross linking with formaldehyde hippocampal chromatin was extracted and fragmented by sonication by restriction digestion or sonication as previously described (12) and incubated in the presence of a mouse IgG antibody (Upstate), anti-CTCF (Abcam) or anti-Jun antibody (Sigma). Chromatin-antibody complexes were recovered using pre-blocked (1mg/ml BSA and 1ug/ml salmon sperm DNA) Dynabeads Protein A (Life technologies). Purified DNA was analysed by quantitative PCR using Roche Lightcycler480 using SYBR green reagents (Roche). The following primers were designed to assess luciferase plasmid concentrations to normalise transfection efficiencies (LucChIP for;CTTGCAGTTCTTCATGCCCG, LucChIP rev; CTCACGAATACGACGGTGGG) and to assess immunoprecipitation of human ECR1T or ECR1C (hECR1 for; TTAATGGCAGCTACATCCCC, hECR1 rev; TCATGGCAGGAAAACTGCTC).

### Generation of gRNA molecules by a novel annealed oligo template (AOT) method

Single guide RNA (sgRNA) molecules were designed to disrupt the mouse ECR1 enhancer (mECR1; Fig. 1) using the optimised CRISPR design tool (http://CRISPR.mit.edu/). To remove the need for cloning of the guide RNA template into a plasmid as previously described (13) we devised a cloning-free technique for the production of sgRNA. This involved the annealing and PCR amplification of two oligonucleotides: the first oligonucleotide (80mer; 5’-AAAAAAAGCACCGACTCGGTGCCACTTTTTCAAGTTGATAACGGACTAGCCTTATTTTAA*CTTGCTATTTCTAGCTCTAA*-3’), represented the sgRNA scaffold and acted as a generic oligonucleotide for the generation of all subsequent templates; the second oligonucleotide, (63mer; 5’-<u>TTAATACGACTCACTATA</u>GGNNNNNNNNNNNNNNNNNNNGTTTT*AGAGCTAGAAATAGCAAG* - 3’), where “N” denotes the predicted guide sequence (5’sgRNA; ATAGTAAAAGAACGATTGAA and 3’sgRNA; CTCTCCATGCAAAAATAAGG) underlined sequence represents the T7 promoter and italic sequence represents sequence complimentary to the template scaffold oligo, acted as the targeted sequence to the region of interest. These oligonucleotides were amplified using 30 cycles of a standard PCR reaction (92 °C, 30 secs, 60 °C; 30 secs, 72 °C 30 secs) to produce a 122 bp double strand sgRNA template. 100 ng of this template was used to produce sgRNA using a mMESSAGE mMACHINE T7 *in-vitro* transcription kit (Ambion) described in the manufacturer’s instructions and purified using a Megaclear kit (Ambion) with modifications by Harms et al (2014).

**Figure 1.**
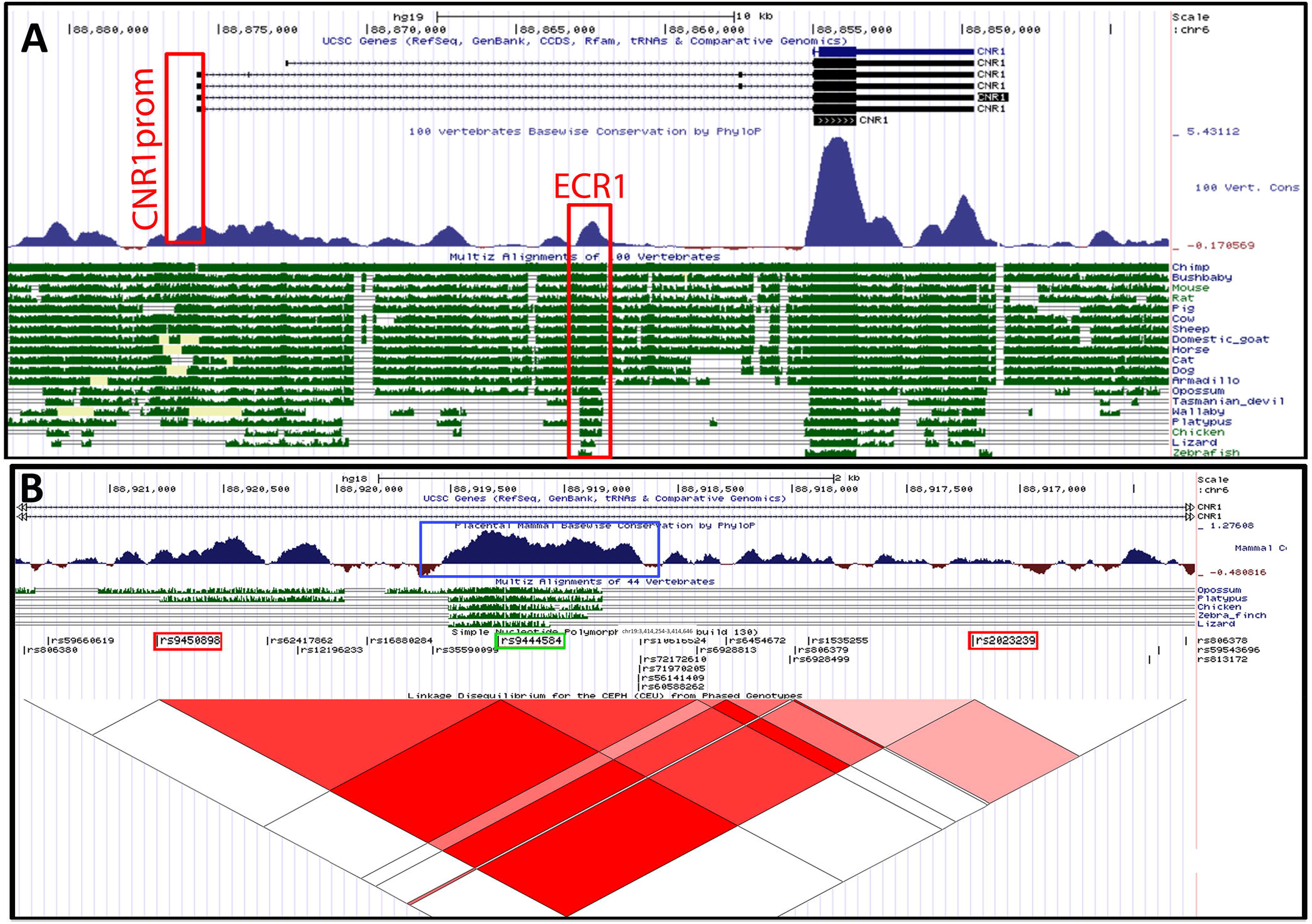
Bioinformatic analysis of the human CNR1 locus using the UCSC genome browser. **A**. Genomic view of 22kb of genomic DNA surrounding the CNR1 locus showing exonic (black bars) and intronic (black lines with chevrons) structure of the gene. Levels of vertebrate conservation (100 vertebrates) is demonstrated using blue peaks and depth of conservation (green bars) spanning 400 million years (fish-human divergence) is displayed in green below. The sequences representing the CNR1 promoter (CNR1prom) and ECR1 are highlighted by red boxes. **B**. Sequence analysis within intron 2 of the human CNR1 locus showing degrees of linkage disequilibrium between disease associated SNPs (rs2023239 and rs9450898; highlighted by red boxes) and rs9444584 (Green box). Deep red triangles indicate linkage disequilibrium expressed as r^2^>0.95 and pink r^2^> 0.9.

### Production of transgenic and genome edited mice

DNA (2ng/μl, linearized plasmid) or RNA (10 ng/μl each gRNA and CAS9 mRNA) were microinjected into the nucleus (DNA) or cytoplasm (RNA) of 1-cell C57/BL6 embryos as described (13). Surviving two-cell embryos were introduced into host CD1 mothers using oviduct transfer as previously described (14). Once weaned, ear-clip biopsies were recovered and analysed by PCR using ECR1 flanking primers (mECR1 test for; TGTGTGCAGAGAGGGGAGAC; mECR1 test rev; CTTTAGGAGTGGACAAGGGGTC) as previously described (13).

### Ethanol intake studies

All animal studies were performed in full accordance with UK Home Office guidelines. Male and female homozygous wildtype and ECR1KO age matched littermates were single housed under standard laboratory conditions (12 h light/12 h dark cycle), with food and water available ad libitum. Oral ethanol self-administration and preference were examined using a two-bottle choice protocol as described previously (15). Mice were housed in TSE home cage systems (TSE, Bad Homburg, Germany) that record liquid intake automatically using weight sensors attached to suspended liquid containers. Initially mice were group housed (~4 / cage) and habituated for 5 days to allow for adaptation to the monitored bottles, then they were single housed for 3 days prior to introduction of the ethanol solution. Intake of water and 10% ethanol solution (both containing 0.2% saccharine to normalise preference to ethanol) were monitored at 60 min intervals over a 3-week period and recorded by TSE PhenoMaster Software.

### Hypothermia response studies

Win55,212-2 was dissolved to 1mg/ml in ethanol and mixed with 2mg/ml tween 80 in ethanol in a 1:1 ratio. Ethanol was evaporated off and the drug/tween 80 mixture was reconstituted in 0.9 % saline. Homozygous wildtype and ECR1KO mice were injected i.p. with 5mg/kg of Win55,212-2 and core temperature measurements taken at 0 minutes, 30 minutes and 60 minutes using a rectal thermometer.

### Data analysis

Statistical significance of data sets was analysed using either a one-way analysis of variance (ANOVA) with Bonferroni post hoc tests or using one tailed or two tailed unpaired parametric Student t-test using GraphPad PRISM version 5.02 (GraphPad Software, La Jolla, CA, USA).

## Results

### CNR1prom supports reporter gene expression in hippocampus and hypothalamic cells

Luciferase reporter studies involving magnetofection of reporters into primary cells demonstrate that the CNR1 promoter(9) (CNR1prom; Fig 1A) is significantly more active in primary hypothalamic and hippocampal cell cultures than the empty luciferase vector. Further analysis of the CNR1prom using a LacZ reporter (pCNR1prom-LacZ; Figure 2A) in 2 separate transgenic mouse lines showed that LacZ gene expression (Fig 2D) did not reflect the expression of the endogenous CNR1 gene in the hippocampus (Fig2 E). However, in both pCNR1prom-LacZ transgenic lines, significant evidence of CNR1 promoter activity was found in the hypothalamus and amygdala that reflected expression of the endogenous gene (Fig 2F and G).

**Figure 2.**
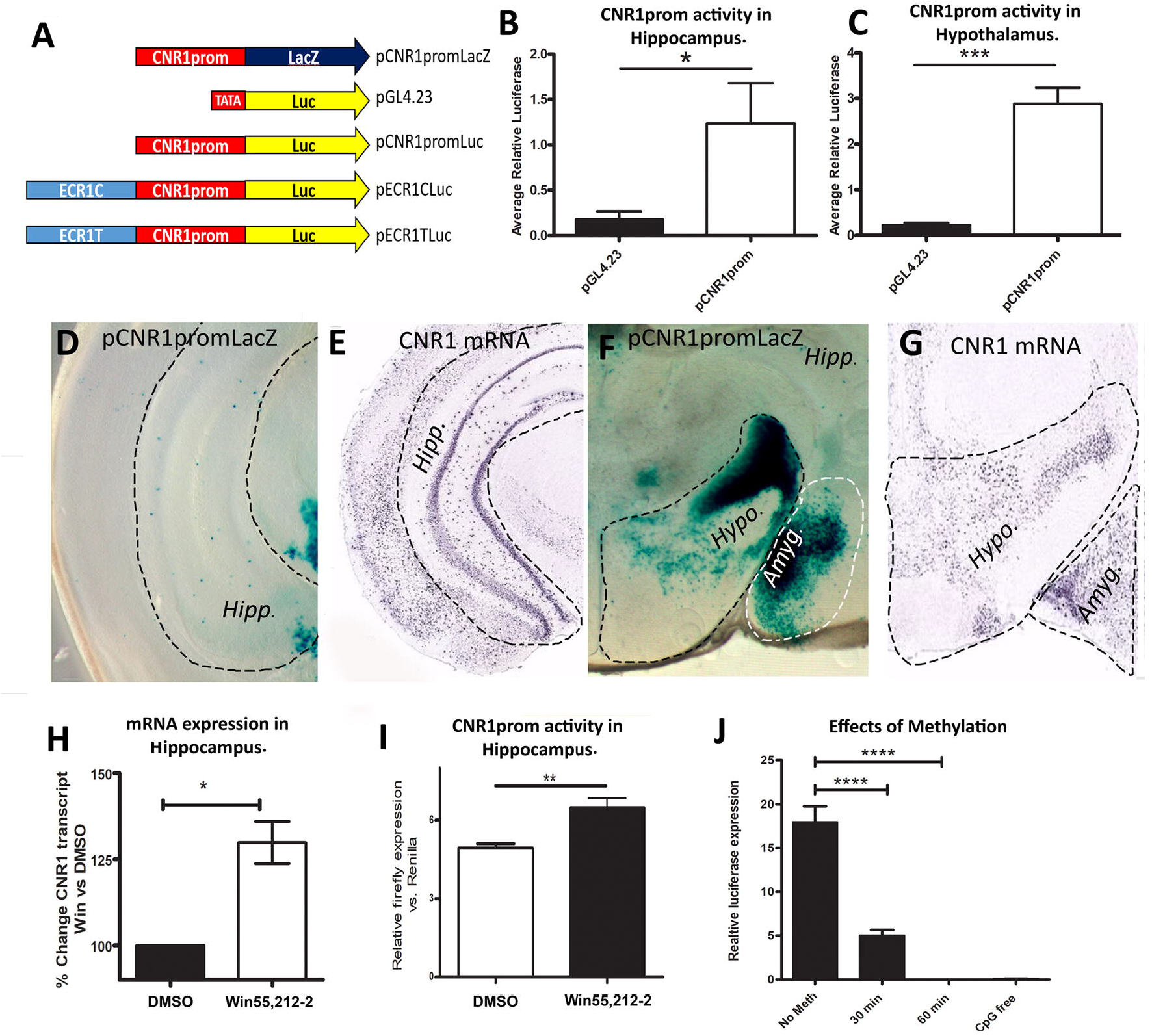
**A**. Diagrammatic representation of the reporter constructs used in the current study demonstrating the relative positions of the ECR1 element (light blue) promoter sequences (red) and reporter genes (blue; β-galactosidase and yellow; firefly luciferase, not to scale). **B and C** results of dual luciferase studies undertaken on primary rat hippocampal (**B**) or hypothalamic cell cultures (**C**) magnetofected with either the empty luciferase vector (pGL4.23) or pCNR1prom-luc (pCNR1prom) luciferase reporter constructs co-transfected and normalised with pGL4.74 (renilla luciferase) (n=14, *;p<0.05, ***;p<0.005). **D** and **F** vibratome sections of tissue derived from mice transgenic for the CNR1prom-lacZ construct after staining with X-gal showing transgene expression in **D**, hippocampus (Hipp.); **F** hypothalamus (Hypo.) and amygdala (Amyg.). **E** and **G** RNA in-situ hybridisation on tissue sections of wild type mice (Allen brain atlas) demonstrating the distribution of CNR1 mRNA within the hippocampus (Hipp.), Hypothalamus (Hypo.) and amygdala (Amyg.). **H**. QrtPCR analysis of rat CNR1 expression in total RNA derived from primary hippocampal neurones treated either DMSO or with 100nM of the CB_1_ agonist Win55,212-2 for 16 hours and normalised against expression of Nono (n=5, *; p<0.05). I. Dual luciferase analysis of primary hippocampal cell cultures magnetofected with pCNR1prom-Luc (plus pGL4.74) and treated with DMSO (control), or Win55,212-2 (n=6, **; p<0.01). **J**. dual luciferase analysis of primary hippocampal cells magnetofected with the pCpG-free-CNR1prom construct that had been exposed to M.SssI for various lengths of time (X axis) (n=6, error bars = SDM, ****; p<0.001).

### CNR1prom responds to activation of the CB_1_ receptor

To confirm previous observations that expression of the CNR1 gene was affected by stimulation of CB1 (16, 17) we treated primary hippocampal neurones with 100nM of the CB_1_ agonist Win55,212-2 (18) or Vehicle (DMSO). After incubation with Win55,212-2, total mRNA was extracted and levels of CNR1 mRNA analysed by QrtPCR to show a significant (15-30%) increase in CNR1 mRNA expression (figure 2H). Furthermore, transfection of CNR1prom-Luc into primary hippocampal cells, and treatment with Win55,212-2 increased the activity of CNR1prom-luc suggesting a role for CNR1prom in CB_1_ autoregulation (Fig 2I).

### CNR1prom is strongly repressed by CpG-methylation in hippocampal cells

To determine the effects of CpG methylation on CNR1 activity we first cloned CNR1prom into a CpG-free luciferase vector that lacks any CpG dinucleotides and exposed these plasmids to the enzyme M.SssI; that specifically methylates at CpG dinucleotides, for varying levels of time up to one hour and transfected these plasmids into primary hippocampal cells. We observed that CpG-methylation strongly repressed CNR1prom activity suggesting that CNR1prom is very susceptible to this environmentally modulated process in hippocampal cells (Fig 2J).

### A highly conserved polymorphic sequence within intron 2 of the CNR1 locus influences CNR1 gene expression in hippocampus and hypothalamus

Intron 2 of the human CNR1 gene contains a 3kb linkage disequilibrium block (LD block) that consists of 17 polymorphisms two of which; rs2023239 and rs9450898 highlighted by red boxes in Fig 1B, are associated with addictive behaviours (19), depression (20), psychosis (21), reduced hippocampal volume in cannabis abuse (22), nicotine addiction (23), obesity (24) and alcohol abuse (25, 26). Because this LD block was non-coding we used comparative genomics to identify functional sequences based on the hypothesis that high (sequence-sequence) and deep (conservation through evolution) conservation reflects functionality (27, 28). We identified a sequence that demonstrated high levels of conservation in all higher vertebrates (Fig 1A and B) and which contained a SNP (rs9444584) in high LD with both rs2023239 and rs9450898 (Fig 1B)(11). To determine the possible function of this sequence we used CRISPR genome editing to disrupt the most highly conserved core region of ECR1 from the mouse genome (Fig 3A) by microinjecting single guide RNA (gRNA; gRNAECR1a and gRNAECR1b; Fig 3A) molecules, designed to disrupt the core region of ECR1, centred on a sequence homologous to that containing the human rs9444584 polymorphism (Fig 3A), together with CAS9 mRNA into the cytoplasm of fertilised single cell mouse embryos. We derived 3 lines containing a deletion that closely matched the PAM sequence of our guides and were unable to detect off target effects (29). These deletions were associated with a 17% reduction in CNR1 mRNA in the hippocampus of these lines (Fig 3B). Although there was some evidence of reduced expression in the hypothalamus it failed to reach significance (Fig 3C). These observations suggest a role for ECR1 as a tissue specific regulatory sequence with greater relevance for CNR1 expression in the hippocampus.

**Figure 3.**
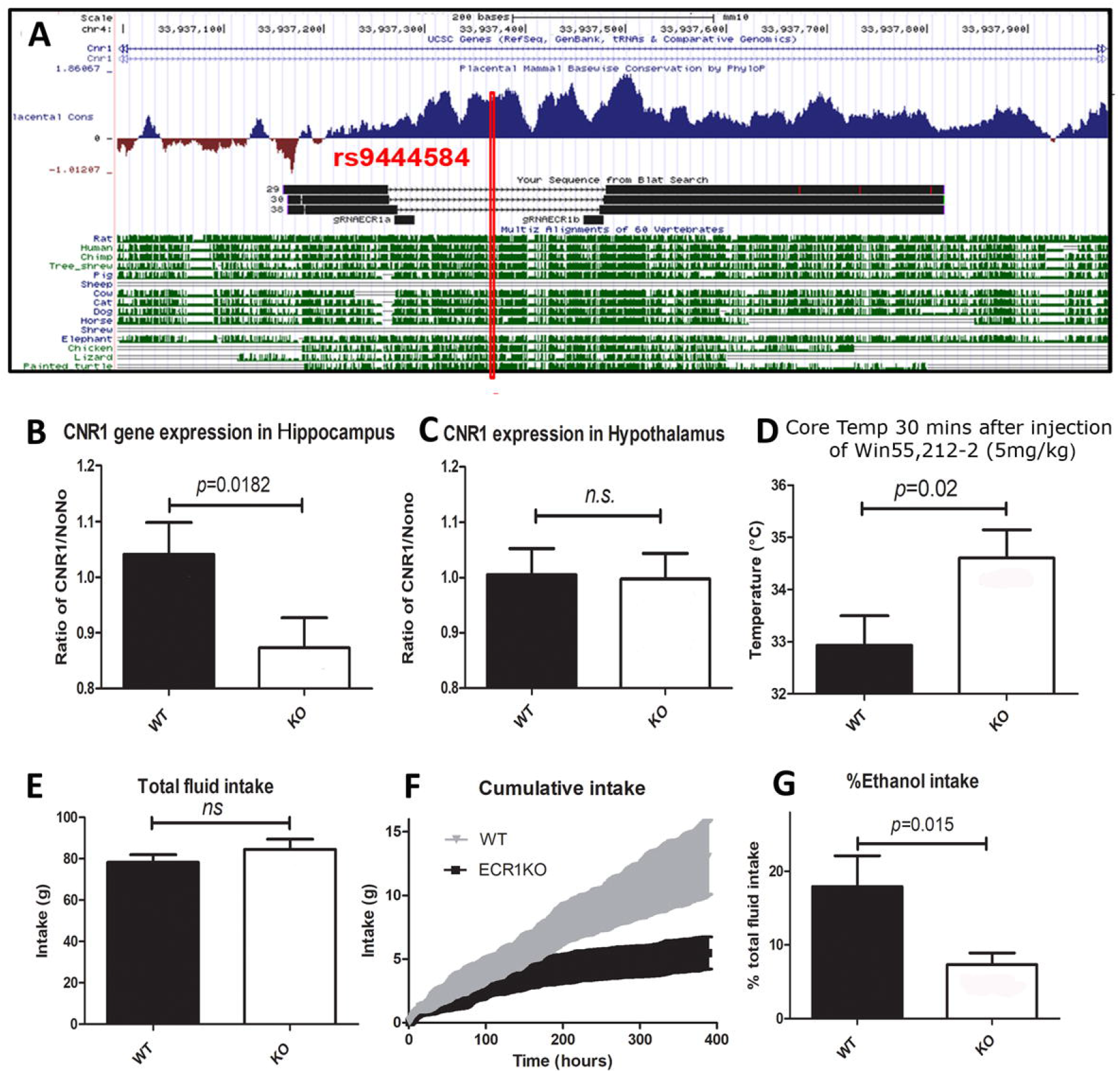
**A** Sequence analysis of the mouse ECR1 enhancer region showing levels of mammalian conservation (blue peaks) and depth of conservation (green bars). Long black bars (29, 30, 38) reflect the alignment of sequences derived from PCR products amplified from DNA of CRISPR genome edited mice. Deletions are indicated by thin lines covered in chevrons. Short black bars (gRNAECR1a and gRNAECR1b) reflect the regions targeted by the guide RNA used to target the ECR1 locus. The red line highlights where the rs9444584 polymorphisms within the ECR1 enhancer lies within the equivalent human locus. **B and C** QrtPCR analysis of mouse CNR1 expression in total RNA derived from **B**, hippocampus and **C**, hypothalamus comparing CNR1 expression in wild type (WT) and ECR1KO (KO) animals (n≥19; error bars=SDM, ns= not significant). **D**, comparison of core temperature measurements from wild type (WT) and ECR1KO (KO) animals taken 30 minutes after intraperitoneal injection with 5mg/kg of Win55,212-2 (n≥17; error bars= SDM). **E**, Comparison of total fluid intake over 400 hours by wild type (WT) and ECR1KO animals given a choice of drinking water or water and 10% ethanol (n≥11; error bars= SDM, ns=no significance). **F**, a time plot comparing cumulative hourly intake of 10% ethanol between WT (grey triangles) and ECR1KO (black squares) age matched littermates over a 400-hour period (n≥11; error bars= SDM). G, a comparison of total final intake of 10% ethanol comparing wild type and ECR1KO animals (n≥11; displayed within bars, error bars= SDM).

### Disruption of ECR1 reduces the CB_1_ activated hypothermia response and reduces ethanol intake

To determine the effects of ECR1 disruption on CB_1_ response we injected adult male and female ECR1KO and wild-type mice with Win55,212-2 (5mg/kg; i.p) and observed that, 30 minutes after injection, ECR1KO mice displayed a significantly reduced hypothermia response compared to wild types (WT) consistent with a reduction in CB_1_ receptor expression (Fig 3D). Because mice lacking CB_1_ also have reduced ethanol intake (15) and the haplotype block harbouring the rs9444584 polymorphism is associated with increased susceptibility to alcohol abuse (26) we explored the effects of the ECR1 knockout on the ethanol intake of ECR1KO mice in comparison to wild type littermates. We observed that, despite having the same total fluid intake (Fig 3E), ECR1KO mice drank significantly less 10% ethanol than WT animals (Fig3F and G).

### Allelic variants of the human ECR1 enhancer differentially regulate the activity of CNR1prom

To explore the tissue specificity of interaction of ECR1 with CNR1prom (30) we cloned allelic variants of ECR1 into the CNR1prom-luc reporter construct (Fig 2A) and magnetofected these constructs into rat hippocampal and hypothalamic primary cell cultures. We found that, although the C-allele did not increase CNR1prom activity in hypothalamus, we observed strong enhancement of CNR1prom activity in hippocampal cells (Fig 4A) demonstrating evidence of enhancer-promoter specificity in these cells. However, the T-allele failed to enhance activity of CNR1prom in hippocampal neurones and repressed its activity in hypothalamus (Fig 4A).

**Figure 4.**
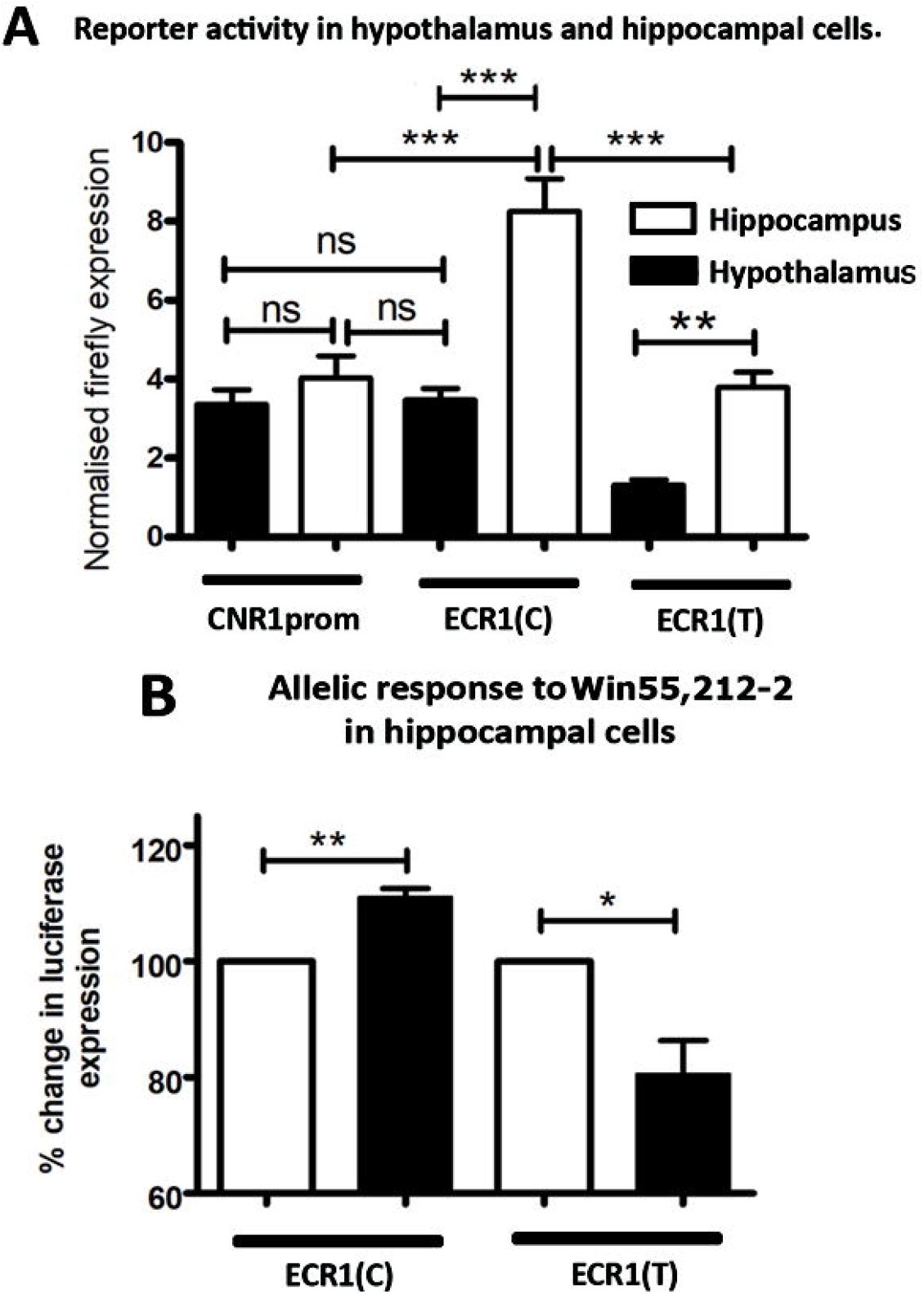
**A** Dual luciferase analysis comparing the relative activity of allelic variant of the ECR1 sequence in primary hypothalamic and hippocampal cells magnetofected with pCNR1prom-Luc (pCNR1prom), pECR1Cluc (pECR1(C)) or pECR1Tluc (pECR1(T) see fig 2A), (n=14, error bars = SDM, *; p<0.05, **; p< 0.01; ***; p< 0.005, ns=no significance). B dual luciferase analysis of primary hippocampal cells magnetofected with pCNR1prom-Luc (pCNR1prom), pECR1(C)luc (pECR1(C)) or pECR1(T)luc (pECR1(T)) constructs and treated with Win55,212-2 (100nM) (n=6, error bars = SDM, *; p<0.05, **; p< 0.01, ns=no significance).

### The ECR1C allele supports the auto-regulatory properties of CNR1prom

Reporter constructs containing allelic variants of the ECR1 enhancer and CNR1prom were magnetofected into hippocampal primary cell cultures which were then treated with Win55,212-2 (100nM). Dual luciferase analysis of these cells demonstrated that incubation of cells with Win55,212-2 significantly increased luciferase expression in the presence of the ECR1(C) allele (Fig 4B). However, we observed a significant reduction in the response of CNR1prom to Win55,212-2 treatment in the presence of ECR1(T) suggesting that ECR1T does not support autoregulation of CNR1prom in hippocampal cells (Fig4B).

### ECR1C interacts with higher affinity to AP-1 transcription factor than ECR1T

Bioinformatic analysis (RegSNP) of the ECR1 locus predicts that the AP-1 transcription factor (a dimer of cFOS and cJUN) binds to ECR1(C) with high affinity but with reduced affinity to ECR1(T). This is an interesting prediction as cFOS and CNR1 are co-expressed in the hippocampus (Fig 5A and B) and several studies have demonstrated that an upregulation of CB1 activity also increases activation of cFOS (31, 32). To explore this prediction, we magnetofected either the ECR1CLuc or ECR1TLuc plasmids (Fig 2A) into primary rat hippocampal cell cultures. After 48 hours chromatin was extracted and fragmented using restriction digestion or sonication (Fig5C and supplementary data). Quantitative PCR of extracted chromatin against the luciferase gene indicated equal transfection efficiencies and were used to normalise transfection. This chromatin was then incubated with antisera against the AP-1 or CTCF proteins. After recovery of specific antibody-protein-DNA complexes QPCR specific for human ECR1 was used to determine comparative levels of human ECR1 DNA immunoprecipitation. Despite using two different genome fragmentation techniques we consistently observed a higher signal from primary cells transfected with ECR1CLuc compared to those transfected with ECR1TLuc suggesting an increased affinity for AP-1 to the C-allele (Fig 5C and supplementary data). Intriguingly, we also observed increased binding of CTCF to the T-allele of ECR1 (Supplementary data).

**Figure 5.**
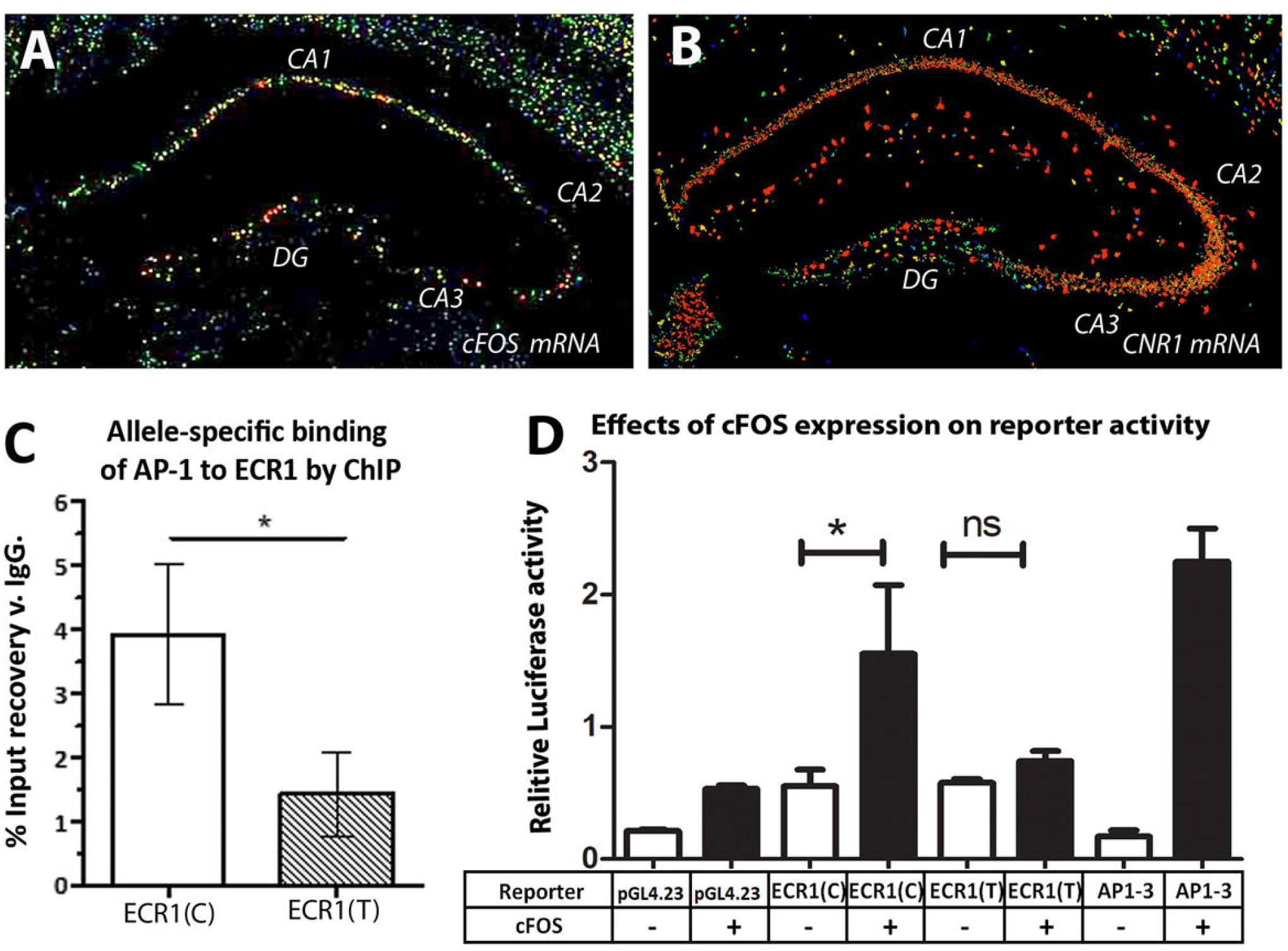
**A and B** *in-situ* hybridisation (Allen brain atlas) of a coronal section of mouse midbrain comparing localisation of **(A)** cFOS and **(B)** CNR1 mRNA within different regions of the hippocampus (CA1, CA2, CA3 and Dentate Gyrus (DG)). **C**. Combined data from qPCR analysis of DNA recovered following chromatin immunoprecipitation (ChIP) of fragmented (restriction digested or sonicated) chromatin derived from primary rat hippocampal cells transfected with pECR1C-luc (pECR1(C)) or pECR1T-luc (pECR1(T)) constructs (See supplementary data for individual experiments). ChIP was undertaken using antisera against AP-1 and QPCR was designed to amplify human ECR1 DNA. (n=4, error bars = SDM, *; p<0.05). **D**. results of a co-transfection study in SH-SY5Y cells where different reporter constructs (pGL4.23; empty luciferase reporter, ECR(C) and ECR1(T); luciferase reporter plasmids containing either the C-allele of T-allele variant of ECR1 respectively, have been co-transfected with either the empty expression vector (pcDNA3.1) or an expression vector expressing the cFOS protein. AP1-3; pAP1-3 positive control plasmids responsive to AP-1 expression. All transfections were normalised against renilla luciferase (pGL4.74) and possible trans effects were negated by normalising against cells transfected with empty vector. (n=3, error bars =SDM, n.s.; not significant, *; p<0.05)

To determine whether the increased binding affinity of AP-1 to ECR1(C) was reflected in enhancer activity we carried out co-transfection analysis in a human neuroblastoma cell line with luciferase reporters containing the C-allele and T-allele of ECR1 in combination with an expression vector expressing the cFOS protein (pcDNA- -FosWT). We observed that co-expression of cFOS with a luciferase reporter containing a promoter known to respond strongly to AP-1 (pAP1-3) was highly up-regulated (Figure 5D). We also observed a significant increase in expression of luciferase in cells co-transfected with ECR1CLuc and pcDNA-cFOS. Consistent with our previous primary cell transfection and ChIP data we found no significant upregulation in activity of ECR1(T) in the presence of cFOS expression (Fig5D).

## Discussion

Although targeting the CB1 receptor has clear therapeutic potential the beneficial effects of cannabinoid drug treatments are not universal (7). Thus, understanding the pharmacogenetics of the CNR1 locus within the human population will be essential to the therapeutic development of the cannabinoids. Because there is evidence that the majority of functional genomic variations occurs outside of coding regions (33) and the CNR1 coding region does not contain polymorphisms that occur in greater than 1% of the population, the current study sought to better understand the noncoding genetic and epigenetic influences controlling the tissue specific expression of the CNR1 gene as possible contributory factors in the pharmacogenomics of cannabinoid response.

We first identified that CNR1prom (9) was active in different primary cell types as well as in specific regions of the brain in transgenic mice expressing a LacZ reporter construct driven by the human CNR1 promoter. In addition, our primary hippocampal cell-based studies support previous observations in T-cells and liver (16, 17) that CNR1 is under autocrine control and we demonstrate that the CNR1 promoter is involved. Moreover, given the known role of CpG-methylation in modulating CNR1 expression (34, 35) our observed sensitivity of CNR1prom to methylation may have important ramifications for understanding the mechanistic effects of environmental factors, such as early life stress, disease or deprivation, on cannabinoid response.

Disrupting the ECR1 enhancer element using CRISPR genome editing demonstrated that ECR1 is important for maintaining normal levels of CNR1 expression within the hippocampus. This experiment also permitted *in-vivo* behavioural studies to determine the effects of ECR1 disruption on core body temperature, following CB1 activation, and ethanol intake. This last observation is interesting as human ECR1 contains the rs9444584 polymorphism; part of a haplotype block that has been implicated in reduced CNR1 expression (9) and increasing susceptibility to alcohol abuse (26). Thus, in addition to its possible role in stratification of drug response, further functional analysis of this polymorphism in vivo and in the clinic may reveal important insights into the causes of alcohol abuse.

From the perspective of tissue specific regulation our observations suggest that ECR1 may play a more substantive role in supporting expression of CNR1 in the hippocampus than the hypothalamus implying that SNPs within ECR1 would have a greater effect in the hippocampus. As we have shown that the C-allele of ECR1 is more active in hippocampal cells, these observations suggest that the human specific C-allele of ECR1 may drive higher expression of CNR1 in hippocampus consistent with previous observations (9). Considering the known neuroprotective role of CB1 activation against the effects of stress in the hippocampus (36) it has not escaped our attention that the C-allele may play a role in protecting the hippocampus against the depressive and anxiety inducing effects of stress. In this context, and from the perspective of adaptive evolution, it is interesting that the C-allele has undergone positive selection in European and Asian populations where its frequency exceeds 80% compared to the ancestral T-allele. Although bottleneck effects might account for this difference, we cannot rule out the possibility that selection for an allele which drives higher levels of CB1 receptor in the hippocampus may have been beneficial to early human populations by resisting the anxiety and depressive effects of stress.

These observed differences in the activity of the C and T-alleles of ECR1 in the hippocampus may also influence the observed effects of CB_1_ targeted therapeutics such as the anti-obesity drug rimonabant in humans. Rimonabant’s main mode of action is through antagonism/inverse agonism of the CB_1_ receptor in the hypothalamus, which leads to a reduction in appetite. Rimonabant was withdrawn from the market as it was linked to an increase in depressed and suicidal feelings in 26% of patients (6). If we consider what is known about the neuroprotective role of CB_1_ against stress in the hippocampus, the increase in anxiety and depression in these individuals is not surprising as the appetite reducing antagonism of CB_1_ in the hypothalamus would be accompanied by antagonism of CB_1_ in the hippocampus. Indeed, reports of increased anxiety (37) and depression-like (38) behaviours have been reported in rodents, who harbour the ancestoral T-allele of ECR1, following chronic administration of rimonabant. What is probably more noteworthy is the observation that 74% of patients did not experience anxiety/depression like symptoms and responded positively to rimonabant. Based on our observations, we propose that the human specific C-allele of ECR1 may induce higher levels of CB1 expression in the hippocampus in humans, thus protecting individuals from the anxiolytic and depression forming effects of stress following treatment with rimonabant (2). Our analysis goes on to identify a possible molecular mechanism that may explain differences in the activity of the C and T-alleles of ECR1 based on variable affinity to the AP-1 transcription factor which is known to be expressed in the hippocampus. Using a unique experiment based on ChIP analyses of magnetofected primary hippocampal cells we demonstrated increased affinity for AP-1 to the C-allele of ECR1 (C). To verify this observation, we also showed that expression of cFOS; one of the proteins that forms the AP-1 complex, activates the ECR1(C) enhancer but not ECR1(T) in co-transfected cells. This observation was interesting as several studies have demonstrates that stimulation of CB_1_ induces binding of AP-1 to DNA and expression of cFOS (31, 32). This was particularly evident in the hippocampus where CB1 and cFOS are co-expressed in the CA1, CA3 and dentate gyrus (39). Further bioinformatics analysis predicted binding of the CTCF transcription factor; a known marker of insulator function, with higher affinity to the T-allele, a prediction that we also support using ChIP assays. These observations raise an interesting possibility that the ancestral T-allele of ECR1 would act as a weak enhancer with insulator properties in the hippocampus of most vertebrates. However, the change to the C-allele in humans, that increased binding affinity of AP-1 and increased the enhancer like properties of ECR1 in specific tissues, may have proved advantageous to specific populations faced with new and stressful circumstances thus expanding the frequency of the C-allele in these populations.

Taken together, the evidence discussed above suggests a role for methylation of CNR1prom and allelic variation within ECR1 in modulating the expression of CNR1. Although much remains to be done to conclusively establish a role for these observations in alcohol abuse and drug effects in humans, our unique primary cell-based and in-vivo studies lay the foundation for future studies of the contribution of these genetic and epigenetic factors in the pharmacogenetics of the cannabinoids and the possible effects on human health and disease susceptibility. We believe that the current manuscript not only provides a platform for the further study of the effects of polymorphisms and DNA-methylation on the pharmacogenomics of the cannabinoids but also provides a functional blueprint for understanding the role of the non-coding genome in health and disease.

## Acknowledgements and conflicts of interest

EH was funded by Medical Research Scotland (PhD-719-2013) and GW Pharmaceuticals. AMcE was funded by BBSRC project grant (BB/N017544/1). PB and DW are funded by the Scottish Government Rural and Environment Science and Analytical Services Division to the Rowett Institute. The authors declare no conflicts of interest.

